# The expression of congenital Shoc2 variants induces AKT-dependent feedback activation of the ERK1/2 pathway

**DOI:** 10.1101/2023.12.23.573219

**Authors:** Patricia Wilson, Lina Abdelmoti, Tianyan Gao, Emilia Galperin

## Abstract

The Shoc2 scaffold protein is crucial in transmitting signals within the Epidermal Growth Factor Receptor (EGFR)-mediated Extracellular signal-regulated Kinase (ERK1/2) pathway. While the significance of Shoc2 in this pathway is well-established, the precise mechanisms through which Shoc2 governs signal transmission remain to be fully elucidated. Hereditary mutations in Shoc2 are responsible for Noonan Syndrome with Loose anagen Hair (NSLH). However, due to the absence of known enzymatic activity in Shoc2, directly assessing how these mutations affect its function is challenging. ERK1/2 phosphorylation is used as a primary parameter of Shoc2 function, but the impact of Shoc2 mutants on the pathway activation is unclear. This study investigates how the NSLH-associated Shoc2 variants influence EGFR signals in the context of the ERK1/2 and AKT downstream signaling pathways. We show that when the ERK1/2 pathway is a primary signaling pathway activated downstream of EGFR, Shoc2 variants cannot upregulate ERK1/2 phosphorylation to the level of the WT Shoc2. Yet, when the AKT and ERK1/2 pathways were activated, in cells expressing Shoc2 variants, ERK1/2 phosphorylation was higher than in cells expressing WT Shoc2. We found that, in cells expressing the Shoc2 NSLH mutants, the AKT signaling pathway triggers the PAK activation, followed by phosphorylation and Raf-1/MEK1/2 /ERK1/2 signaling axis activation. Hence, our studies reveal a previously unrecognized feedback regulation downstream of the EGFR and provide evidence for the Shoc2 role as a “gatekeeper” in controlling the selection of downstream effectors within the EGFR signaling network.

## INTRODUCTION

Hereditary variants in the *shoc2* gene cause a developmental disorder termed Noonan Syndrome with loose anagen hair (NSLH) (OMIM #607721) (1–3). Patients harboring *shoc2* mutations present with several congenital deficiencies, including facial dysmorphia (e.g., ocular hypertelorism), cleft palate, cardiac abnormalities, short stature, and other less frequent abnormalities (3–7). Shoc2 *null* mice and zebrafish CRISPR/Cas9 knockouts develop systemic deficiencies resulting in early embryonic lethality, manifesting the biological significance of Shoc2 in vertebrate development, morphogenesis, and the maintenance of various tissues in adult organisms (8–10).

Shoc2 is a non-redundant evolutionary conserved signaling scaffold protein. The best-studied function of Shoc2 is its ability to accelerate signals of the extracellular signal-regulated kinase 1/2 (ERK1/2) pathway (11). Several reports also show that Shoc2 plays a role within the AKT, mTOR, and PI3K signaling pathways (12,13), although these aspects of the Shoc2 function must be better understood. Shoc2 interacts with different protein partners. It tethers RAS and the catalytic subunit of protein phosphatase 1c (PP1c, also known as PP1CA) to accelerate ERK1/2 signals *via* dephosphorylation of inhibitory phospho-S259 of RAF-1 (14).

Shoc2 also recruits several proteins of the ubiquitin system: HUWE1, VCP, USP7, PSMC5, and FBXW7 (15–19) to form machinery that modulates the ability of Shoc2 to accelerate the ERK1/2 signals and facilitate cross-talk with other signaling pathways (20). Due to the critical role of Shoc2 in transmitting ERK1/2 signals, significant effort has been invested in ascertaining the structure of the Shoc2-MRas-PP1C complex and gaining insights into RAF activation. These studies provided substantial insight into the assembly of the Shoc2-MRas-PP1C complex and the potential role of the specific amino residues in the stability of the holoenzyme complex (21–24).

Eight pathogenic *shoc2* variants have been reported in individuals clinically diagnosed with NSLH, and several others can be found in public databases (2,3,7,15,25). The pathogenicity of these variants is commonly established in the context of the Epidermal Growth Factor Receptor (EGFR) activation by examining the levels of ERK1/2 phosphorylation downstream of EGFR (2,7,25–27). However, most studies classifying Shoc2 variants do not consider different signal initiation events (i.e., ligands and levels of active receptors) and the adaptive responses limiting or promoting intracellular signal transduction downstream of EGFR. The work of several labs has shown that specific subsets of signaling cascades downstream of EGFR are activated by different concentrations of EGF. For instance, EGF concentrations that are typically detectable in most human tissues or tumors (∼1-2 ng/ml) (28–32) are sufficient to stimulate phosphorylation of 75–80% of MEK1/2 and ERK1/2 in several types of cultured cells and tumors (33–37). It has also been shown that EGF concentrations ranging from 20–100 ng/ml activate several other signaling pathways downstream of EGFR, including tyrosine phosphorylation of phospholipase C γ1, JAK-STAT3, and PI3K-AKT pathways (33–37).

Here, we examined how the Shoc2 NSLH variants impact intracellular signals downstream of EGFR in the context of the ERK1/2 and AKT signaling pathways. We found that Shoc2 is dispensable for the activation of the AKT pathway. We also found that when the ERK1/2 pathway is the primary signaling cascade activated by EGFR, Shoc2 NSLH mutants fail to enhance ERK1/2 activity to the levels of control cells. Yet, under conditions activating the ERK1/2 and AKT pathways, the levels of ERK1/2 phosphorylation were higher in cells expressing NSLH mutant than those in control cells. Mechanistically, our experiments revealed that in cells lacking Shoc2 or expressing Shoc2 NSLH variants, AKT activation leads to phosphorylation of p21-activated kinase (PAK) followed by phosphorylation of RAF1 and MEK1/2. Together, we demonstrate that Shoc2 preferentially guides EGFR signals through the ERK1/2 pathway. In cells lacking functional Shoc2, the AKT pathway may redirect EGFR signals toward a feedback loop that facilitates ERK1/2 phosphorylation in the Shoc2-independent manner. Our study provides novel insights into the Shoc2 function within the EGFR network and emphasizes the importance of assessing the Shoc2 heritable variants in the context of complex EGFR signaling.

## RESULTS

### EGFR activates the AKT pathway in a Shoc2-independent manner

To understand the role of the Shoc2 scaffold in the crosstalk of the ERK1/2 and AKT signaling pathways, we compared how the Shoc2 loss alters the activation of the ERK1/2 and AKT signaling pathways initiated by different concentrations of EGF. First, we determined that EGF concentrations within the ∼0.2-2 ng/ml range are sufficient to activate close to 75% of phospho-ERK1/2 in Hela cells. We also found that EGF concentrations higher than 2 ng/ml trigger significantly higher levels of EGFR phosphorylation that coincide with a noticeable increase in AKT phosphorylation (**Supp.** Fig. 1). These data align with the findings of the plethora of other studies dissecting the EGFR-ligand relations with respect to activation of downstream pathways (35,36). Hence, in the following experiments, we stimulated cells with either 0.2 ng/ml of EGF or 30 ng/ml EGF, commonly used in Shoc2-related studies (25,27,38).

We then analyzed how the loss of Shoc2 affects the phosphorylation of ERK1/2 and AKT. Here, we utilized Shoc2 CRISPR knockout HeLa cells (Shoc2-KO) and Cos1 cells that have endogenous Shoc2 stably silenced by shRNA (Shoc2-KD). HeLa (Shoc2-KO) and Cos1 (Shoc2-KD) cells were developed and reported previously (3,15). Similar to earlier reports, the striking effect of either Shoc2 knockout or knockdown on ERK1/2 pathway activation was evident in cells stimulated with either 0.2 ng/ml or 30 ng/ml concentrations of EGF (**Fig. 1A, B, D, and E**). Yet, when stimulated with 30 ng/ml of EGF, elevated AKT phosphorylation was readily detectable in cells either expressing endogenous Shoc2 or lacking thereof (up to 2.5-fold change), as measured by phosphorylation of the C-terminal S473 of AKT (**Fig. 1A, D, C, and F**). These results show that Shoc2 is dispensable for activation of the AKT pathway and prompted an additional exploration of the crosstalk between the AKT and the ERK1/2 signaling pathways in the context of the Shoc2 heritable variants.

**Figure 1.**
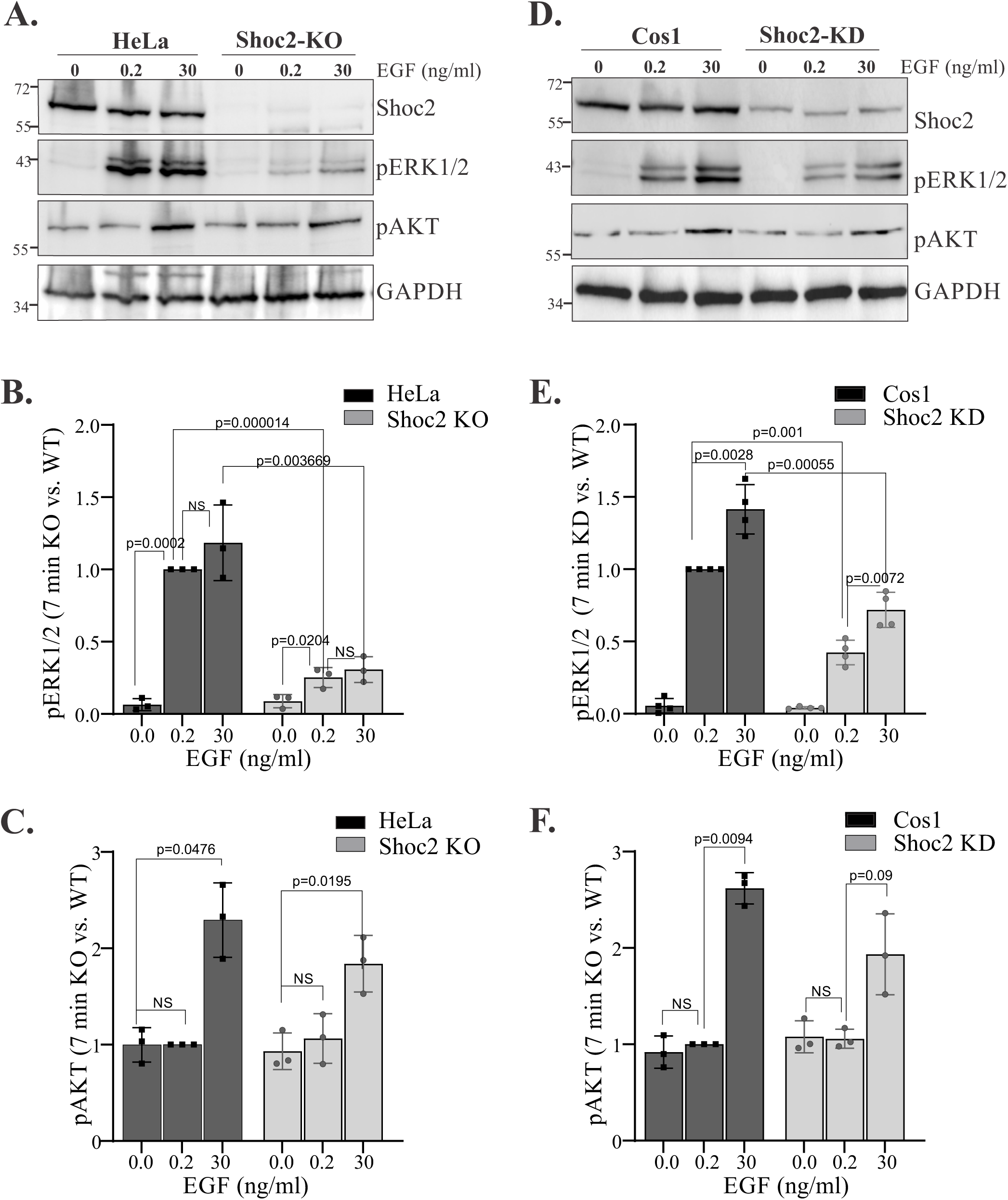
High EGF concentrations activate the AKT pathway in a Shoc2-independent manner. **A.** HeLa and HeLa Shoc2 CRISPR KO cells (Shoc2-KO) were serum-starved for 16 hr and then stimulated with EGF (0.2 or 30 ng/ml) for 7 min at 37°C. Cell lysates were analyzed using anti-pERK1/2, –pAKT, –GAPDH, and –Shoc2 antibodies. **B.** Bars represent the mean amount of pERK1/2 in HeLa-KO cells treated with 0.2 or 30 ng/ml EGF normalized to the total amount of GAPDH in arbitrary units as compared to parental HeLa cells treated with 0.2 ng/ml at 7min ± S.D. (*n*=3) *(p values are indicated on graph,* two-way ANOVA). The results in each panel are representative of those from three independent experiments. NS-no statistical difference was found. **C.** Bars represent the mean amount of pAKT in HeLa-KO cells treated with 0.2 or 30 ng/ml EGF normalized to the total amount of GAPDH in arbitrary units as compared to parental HeLa cells treated with 0.2 ng of EGF at 7min ± S.D. (*n*=3) *(p values are indicated on graph,* two-way ANOVA). The results in each panel are representative of those from three independent experiments. **D.** Cos1 cells constitutively depleted of Shoc2 (Shoc2-KD) were serum-starved for 16 hr and then stimulated with EGF (0.2 or 30 ng/ml) for 7 min. Cell lysates were analyzed using anti-pERK1/2, –pAKT, –GAPDH, and –Shoc2 antibodies. **E.** Bars represent the mean amount of pERK1/2 in Cos1 Shoc2-KD cells treated with 0.2 or 30 ng/ml EGF normalized to the total amount of GAPDH in arbitrary units as compared to parental HeLa cells treated with 0.2 ng/ml at 7min ± S.D. (*n*=4) *(p values are indicated on graph,* two-way ANOVA). The results in each panel are representative of those from four independent experiments. **F.** Bars represent the mean amount of pAKT in Cos1 Shoc2-KD cells treated with 0.2 or 30 ng/ml EGF normalized to the total amount of GAPDH in arbitrary units as compared to parental Cos1 cells treated with 0.2 ng of EGF at 7min ± S.D. (*n*=3) *(p values are indicated on graph,* two-way ANOVA). The results in each panel are representative of those from three independent experiments.

### AKT activation in cells expressing the Shoc2 heritable variants coincides with increased ERK1/2 activation

NSLH is associated with distinct missense mutations in the *shoc2* gene. The Shoc2 (S^2^G) substitution is the most frequent NSLH pathogenic variant (2). Other reported NSLH variants include M^173^I (3) and QH^269/270^HY (3,25). We have previously assessed additional Shoc2 variants reported in individuals exhibiting a striking overlap with the clinical phenotypes observed in NSLH patients (E^89^D, C^238^Y, and L^473^I) using cellular and zebrafish vertebrate models (15). We showed that similarly to Shoc2^S2G^ and Shoc2^QH269/270HY^, the Shoc2^L437I^ variant is likely to be NSLH causative and showed that, unlike Shoc2^WT^, Shoc2 variants Shoc2^S2G^, Shoc2^L437I^ or Shoc2^QH269/270YH^ could not rescue early erythropoietic defects observed in zebrafish Shoc2 knockout larvae (15). These Shoc2 NSLH-related variants were examined here to establish the Shoc2 role in the crosstalk between the AKT and the ERK1/2 signaling pathways.

As noted earlier, the functionality of the Shoc2 heritable variants is often assessed when multiple signaling pathways downstream of EGFR are activated (20-30 ng/ml EGF concentrations). Hence, we compared how different EGF concentrations affect ERK1/2 and AKT phosphorylation in HeLa cells expressing Shoc2 NSLH variants. In these experiments, we utilized the tRFP-fused Shoc2 (Shoc2-tR) (39). To start, we verified that the expression of Shoc2-tR does not affect ERK1/2 phosphorylation in the presence of its endogenous counterpart (**Supp.** Fig. 2A). We then compared how the Shoc2 pathogenic variant S2G (Shoc2^S2G^) affects the phosphorylation of ERK1/2 and AKT in HeLa cells treated with either 0.2 ng/ml or 30 ng/ml (**Fig. 2A**). As expected, phospho-EGFR levels of in cells stimulated with 0.2 ng/ml EGF were lower than in those stimulated with 30 ng/ml EGF (**Fig. 2**). We also found that, in cells expressing the Shoc2^S2G^ variant and treated with 0.2 ng/ml of EGF, phospho-ERK1/2 levels were significantly lower than in cells expressing Shoc2-tR^WT^ (**Fig. 2A, E)**. However, 30 ng/ml of EGF elicited significantly higher ERK1/2 phosphorylation in cells expressing Shoc2^S2G^ than in cells expressing wild-type (WT) Shoc2-tR. Similar observations were made when Shoc2^S2G^ was expressed in Cos1 or 293FT cells (**Supp.** Fig. 2B), recapitulating earlier findings (2,7,25). Interestingly, similar to the results in **Fig. 1A**, differences in phospho-ERK1/2 levels in cells expressing Shoc2^WT^ and treated with low or high EGF concentrations were not dramatic.

**Figure 2.**
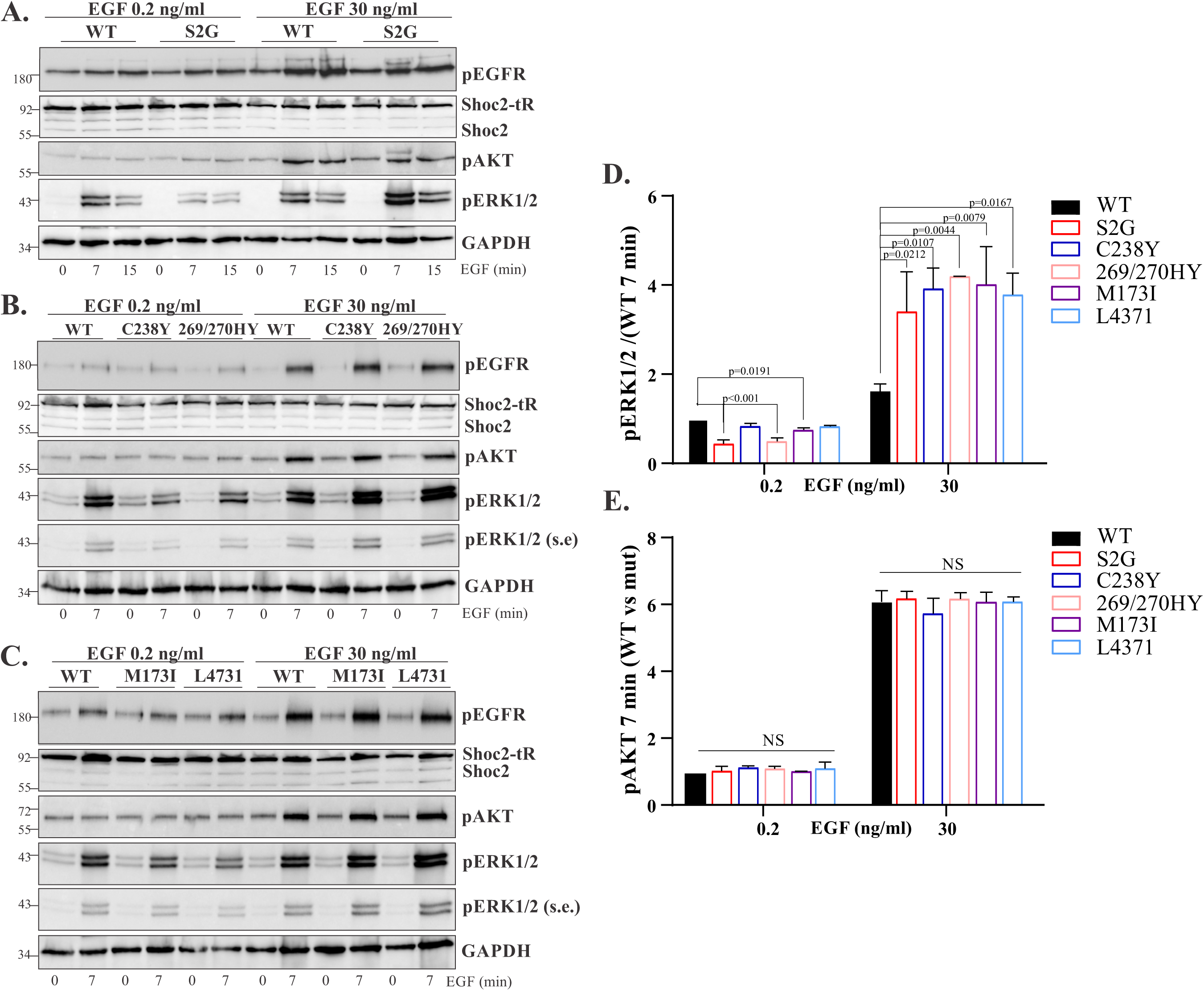
Low and high EGF concentrations differentially activate the ERK1/2 and AKT pathways in cells expressing Shoc2 NSLH mutants. **A.** HeLa cells transiently transfected with the Shoc2-tRFP or Shoc2-tRFP S2G mutant were serum-starved for 16 hr and then stimulated with EGF (0.2 and 30 ng/ml) for 7-or 15-min at 37°C. Cell lysates were analyzed using anti-pEGFR, –pERK1/2, –pAKT, –GAPDH, and –Shoc2 antibodies. **B.** HeLa cells transiently transfected with the Shoc2-tRFP, Shoc2-tRFP (C238Y), or (QH269/270HY) mutants were serum-starved for 16 hr and then stimulated with EGF (0.2 or 30 ng/ml) for 7 min at 37°C. Cell lysates were analyzed using anti-pEGFR, –pERK1/2, –pAKT, – GAPDH, and –Shoc2 antibodies. s.e. – short exposure. **C.** HeLa cells transiently transfected with the Shoc2-tRFP, Shoc2-tRFP (M173I), or (L473I) mutants were serum-starved for 16 hr and then stimulated with EGF (0.2 or 30 ng/ml) for 7 min at 37°C. Cell lysates were analyzed using anti-pEGFR, –pERK1/2, –pAKT, –GAPDH, and –Shoc2 antibodies. s.e. – short exposure. **D.** Bars represent the mean amount of pERK1/2 in cells expressing Shoc2 mutants normalized to the total amount of GAPDH in arbitrary units as compared to cells expressing WT Shoc2-tRFP at 7min ± S.E. (*n*=3) *(p values as shown on graph,* Anova one way). The results in each panel are representative of those from three independent experiments. **E.** Bars represent the mean amount of pAKT in cells expressing Shoc2 mutants normalized to the total amount of GAPDH in arbitrary units as compared to cells expressing WT Shoc2-tRFP at 7min ± S.E. (*n*=3) *(p values as shown on the graph,* Anova one way, NS= non-significant). The results in each panel are representative of those from three independent experiments.

Next, we examined ERK1/2 phosphorylation in cells expressing additional NSLH-associated Shoc2 variants. As in **Fig. 2A**, when stimulated with 0.2 ng/ml of EGF, ERK1/2 phosphorylation was significantly lower in cells expressing Shoc2^C238Y^, Shoc2^QH269/270HY^, Shoc2^M173I^ or Shoc2^L473I^ variants than in cells expressing Shoc2^WT^ (**Fig. 2B-E**). However, when treated with 30 ng/ml of EGF, phospho-ERK1/2 levels were significantly higher in cells expressing Shoc2 NSLH variants than in cells expressing Shoc2^WT^. Consistently, 30 ng/ml EGF triggered increased AKT phosphorylation, readily detectable in cells expressing either WT Shoc2 or the Shoc2 NSLH mutants (**Fig. 2A-F**).

To gain further insight into the Shoc2-mediated ERK1/2 signal transmission, experiments as described above and shown in **Fig. 2** were performed using HeLa cells lacking endogenous Shoc2 (HeLa-KO) (**Fig. 3**). As expected, the levels of phosphorylated EGFR were significantly higher in cells exposed to 30 ng/ml EGF than in cells treated with 0.2 ng/ml EGF. We consistently found that, when treated with 0.2 ng/ml of EGF, the ERK1/2 phosphorylation was significantly lower in cells expressing the Shoc2^S2G^ variant than in those expressing Shoc2^WT^ (**Fig. 3A, D**). However, 30 ng/ml EGF elicited an opposite response and stimulated higher ERK1/2 phosphorylation in cells expressing the Shoc2^S2G^ variant than those expressing Shoc2^WT^. Similar results were obtained when variants Shoc2^QH269/270HY^ and Shoc2^M173I^ were examined (**Fig. 3B-E**). Yet again, only high EGF concentrations triggered a detectable increase in the phosphorylation of AKT (**Fig. 3A, B, C, and F**). In addition, we found that when stimulated with 0.2 ng/ml EGF, ERK1/2 phosphorylation in cells expressing the Shoc2^L473I^ variant was similar to that of cells expressing Shoc2^WT^. However, activation of the AKT pathway (30 ng/ml of EGF) elicited significantly higher levels of phospho-ERK1/2 in cells expressing the Shoc2^L473I^ variant compared to Shoc2^WT^ (**Fig. 3C-E**).

**Figure 3.**
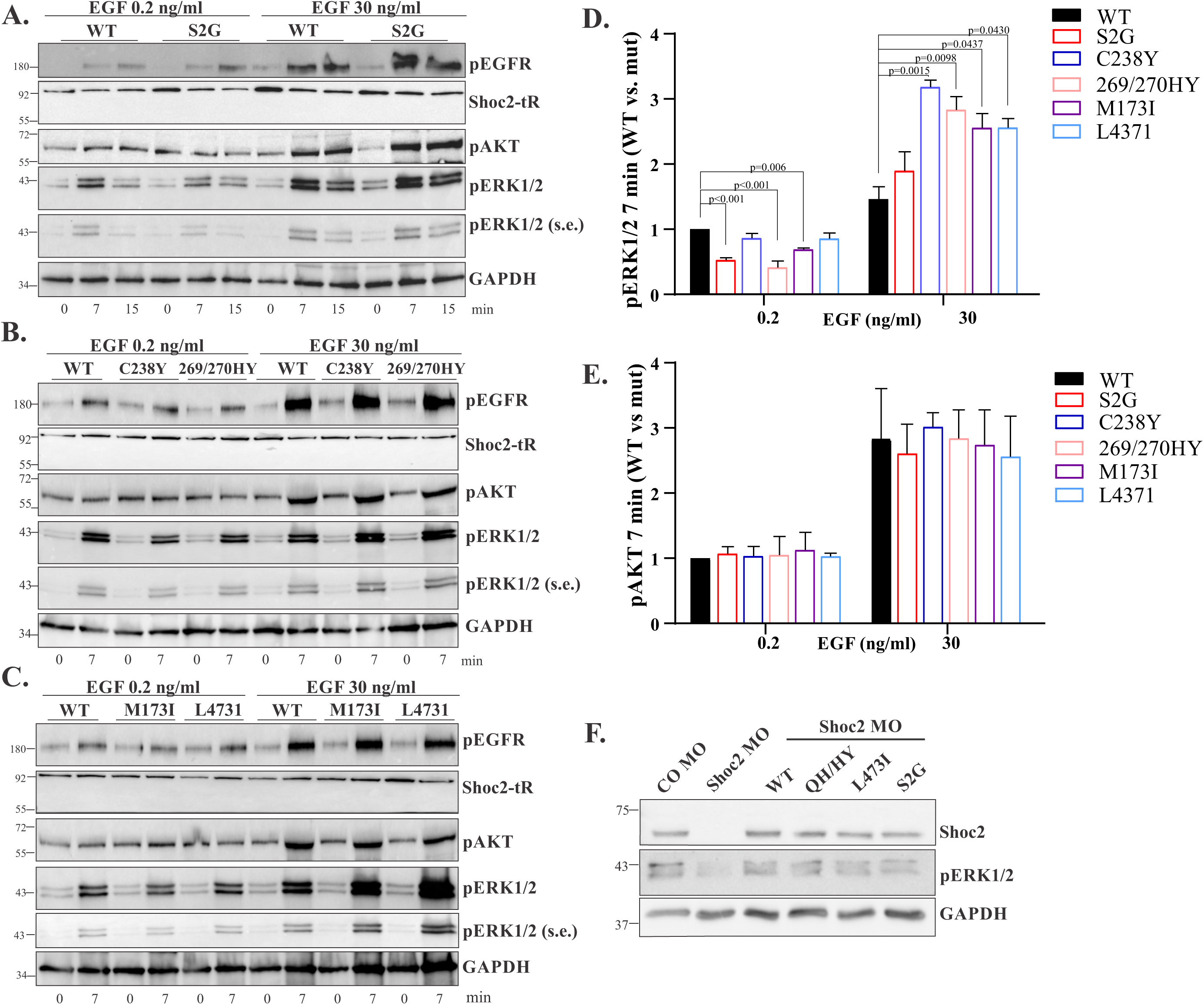
Figure 2. Low and high EGF concentrations differentially activate the ERK1/2 and AKT pathways in cells expressing Shoc2 NSLH mutants in cells lacking endogenous Shoc2. **A.** HeLa-KO cells transiently transfected with the Shoc2-tRFP or Shoc2-tRFP S2G mutant were serum-starved for 16 hr and then stimulated with EGF (0.2 and 30 ng/ml) for 7-or 15-min at 37°C. Cell lysates were analyzed using anti-pEGFR, –pERK1/2, –pAKT, –GAPDH, and –Shoc2 antibodies. s.e. – short exposure. **B.** HeLa cells transiently transfected with the Shoc2-tRFP, Shoc2-tRFP (C238Y), or (QH269/270HY) mutants were serum-starved for 16 hr and then stimulated with EGF (0.2 or 30 ng/ml) for 7 min at 37°C. Cell lysates were analyzed using anti-pEGFR, –pERK1/2, –pAKT, – GAPDH, and –Shoc2 antibodies. s.e. – short exposure. **C.** HeLa-KO cells transiently transfected with the Shoc2-tRFP, Shoc2-tRFP (M173I), or (L473I) mutants were serum-starved for 16 hr and then stimulated with EGF (0.2 or 30 ng/ml) for 7 min at 37°C. Cell lysates were analyzed using anti-pEGFR, –pERK1/2, –pAKT, –GAPDH, and –Shoc2 antibodies. s.e. – short exposure. **D.** Bars represent the mean amount of pERK1/2 in cells expressing Shoc2 mutants normalized to the total amount of GAPDH in arbitrary units as compared to cells expressing WT Shoc2-tRFP at 7min ± S.E. (*n*=3) *(p values as shown on graph,* Anova one way). The results in each panel are representative of those from three independent experiments. **E.** Bars represent the mean amount of pAKT in cells expressing Shoc2 mutants normalized to the total amount of GAPDH in arbitrary units as compared to cells expressing WT Shoc2-tRFP at 7min ± S.E. (*n*=2) *(p values as shown on the graph,* Anova one way, NS= non-significant). The results in each panel are representative of those from three independent experiments. **F.** Embryos injected with shoc2 and control MO and indicated mRNA were harvested for immunoblotting at 52 h post-fertilization. The protein levels were analyzed using specific Shoc2, pERK1/2, and GAPDH antibodies.

We then used the zebrafish vertebrate model to examine our findings in a physiological environment. We compared whether WT Shoc2 and Shoc2 harboring S2G, L437I, and QH269/270YH substitutions can rescue the loss of phospho-ERK1/2 in zebrafish embryos depleted of Shoc2. As we demonstrated previously, embryos injected with shoc2 morpholino (MO) almost completely lacked phospho-ERK1/2 (**Fig. 3F**). Injecting embryos with WT shoc2 zebrafish mRNA (Shoc2^WT^) rescued the shoc2 MO-induced defects in phospho-ERK1/2. We have also observed that levels of phospho-ERK1/2 in embryos injected with shoc2 mRNA harboring Shoc2^S2G^, Shoc2^L437I^, or Shoc2^QH269/270YH^ mutations were close but did not exceed the phospho-ERK1/2 levels detected if embryos injected with Shoc2^WT^ mRNA, supporting our observations in **Fig. 2A-3**. Significantly, these findings prompted us to explore whether the increases in phospho-ERK1/2 levels observed in cells expressing the Shoc2 NSLH variants and treated with 30 ng/ml of EGF stem from the crosstalk between AKT and ERK1/2 signaling pathways.

### The AKT-PAK signaling feedback loop activates ERK1/2 in cells lacking functional Shoc2

To assess the possible contribution of the AKT pathway to the increased levels of ERK1/2 phosphorylation observed in cells treated with 30 ng/ml EGF concentrations, we utilized the selective AKT inhibitor MK2206 (**Fig. 4**). Here, we focused on the subset of the Shoc2 variants, including Shoc2^S2G^, Shoc2^QH269/270HY^, and Shoc2^M173^. HeLa-KO cells transiently expressing WT Shoc2-tRFP or Shoc2 NSLH variants Shoc2^S2G^ (**Fig. 4A**), Shoc2^QH269/270HY^ (**Fig. 4B**), or Shoc2^M173I^ (**Fig. 4C**) were treated with MK2206 (150nM) for 1 hour and then stimulated with 30 ng/ml of EGF. As shown in **Fig. 4A-D**, MK2206 treatment that effectively abrogated AKT phosphorylation also reduced phosphorylation of ERK1/2 in cells expressing Shoc2 inheritable variants (**Fig. 4D**). Importantly, MK2206 treatment had little effect on phospho-ERK1/2 levels in cells expressing WT Shoc2. This data suggested that the AKT pathway likely contributes to the elevated levels of phospho-ERK1/2 in cells expressing Shoc2 NSLH variants (**Fig. 2-4**).

**Figure 4.**
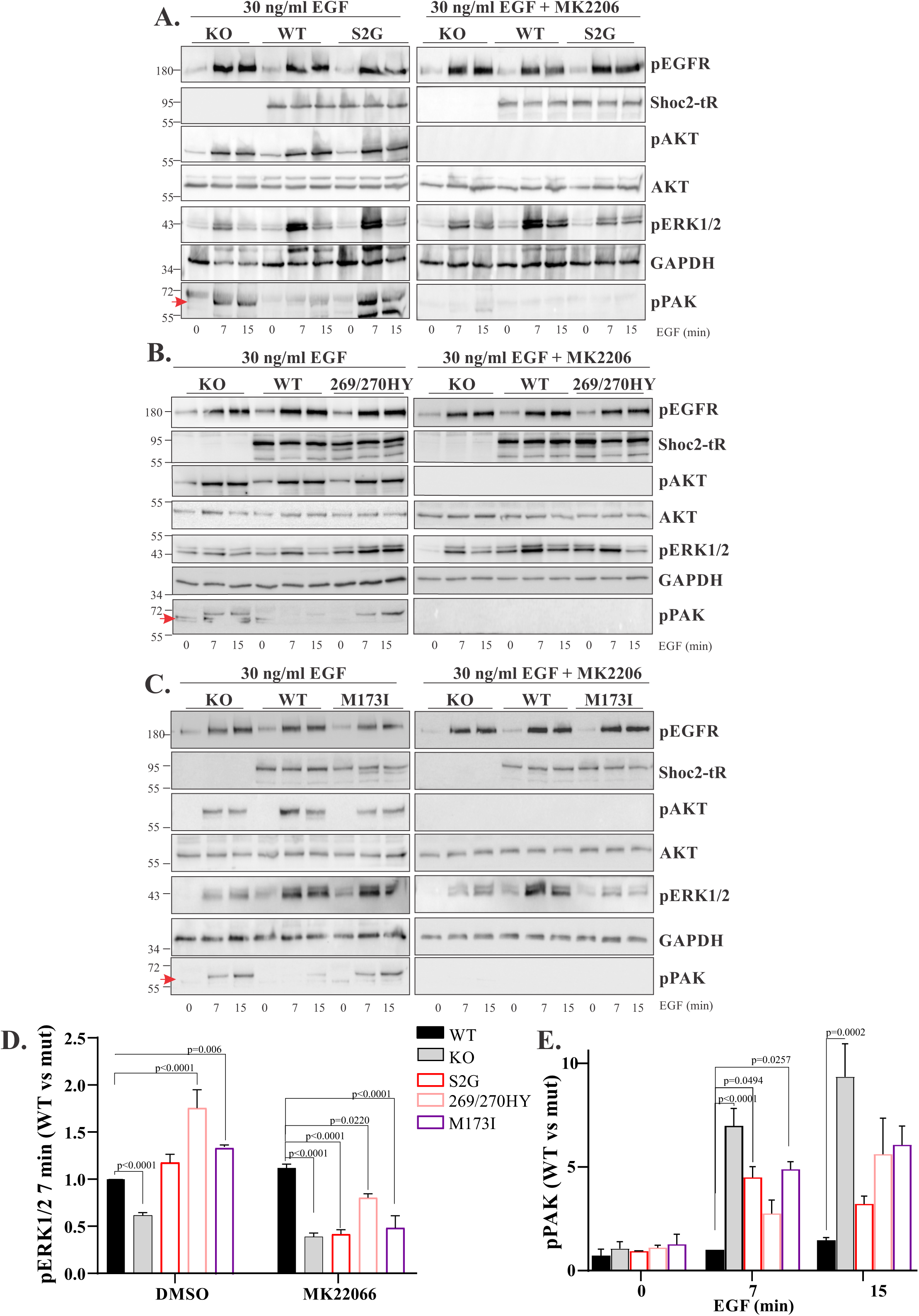
High EGF concentrations activate the AKT feedback loop in a Shoc2-independent manner. **A.** HeLa-KO cells transiently transfected with the Shoc2-tRFP or Shoc2-tRFP (S2G) mutant were serum-starved for 16 hr, treated with DMSO or MK2206 for 1 hour, and then stimulated with EGF (0.2 and 30 ng/ml) for 7-or 15-min at 37°C. Cell lysates were analyzed using anti-pEGFR, –Shoc2, –pERK1/2, –pAKT, –pPAK, AKT, and –GAPDH antibodies. **B.** HeLa cells transiently transfected with the Shoc2-tRFP and Shoc2-tRFP (QH269/270HY) mutant were serum-starved for 16 hr, treated with MK2206 for 1 hour, and then stimulated with EGF (0.2 and 30 ng/ml) for 7-or 15-min at 37°C. Cell lysates were analyzed using anti-pEGFR, –Shoc2, –pERK1/2, –pAKT, –pPAK, AKT, and –GAPDH antibodies. **C.** HeLa-KO cells transiently transfected with the Shoc2-tRFP, Shoc2-tRFP (M173I) mutant were serum-starved for 16 hr, treated with MK2206 for 1 hour, and then stimulated with EGF (0.2 and 30 ng/ml) for 7-or 15-min at 37°C. Cell lysates were analyzed using anti-pEGFR, – Shoc2, –pERK1/2, –pAKT, –pPAK, AKT, and –GAPDH antibodies. **D.** Bars represent the mean amount of pERK1/2 in cells expressing Shoc2 mutants normalized to the total amount of GAPDH in arbitrary units as compared to cells expressing WT Shoc2-tRFP at 7min ± S.E. (*n*=3) *(p values as shown on graph,* Anova one way). The results in each panel are representative of those from three independent experiments. **E.** Bars represent the mean amount of pPAK in cells expressing Shoc2 mutants normalized to the total amount of GAPDH in arbitrary units as compared to cells expressing WT Shoc2-tRFP at 7min ± S.E. (*n*=2) *(p values as shown on graph,* Anova one way). The results in each panel are representative of those from three independent experiments. Red arrows denote the pPAK chemiluminescence signal used for quantification.

ERK1/2 and AKT have broad crosstalk between the two pathways (40). AKT can directly inhibit the ERK1/2 pathway *via* phosphorylation of Ser259 on RAF-1 or Ser364 and Ser428 on B-Raf, thus inhibiting Raf activity (41,42). The AKT signaling cascade also can activate the ERK1/2 pathway through the Rac-PAK-mediated mechanism (43). For instance, PAK1 has been shown to stimulate ERK1/2 through the phosphorylation of RAF-1 and the formation of the MEK1-ERK1/2 complex (44–46). Thus, to obtain additional insight into the mechanisms of AKT-mediated ERK1/2 phosphorylation, we examined levels of phospho-PAK. As shown in **Fig. 4A-C and E**, activation of AKT coincided with the phosphorylation of PAK in cells lacking Shoc2 or in cells expressing the NSLH variants. Of note, in cells expressing WT Shoc2, AKT activation did not coincide with PAK phosphorylation. PAK phosphorylation was abrogated in cells treated MK2206, indicating that high EGF concentrations trigger the EGFR-AKT-PAK axis in cells lacking functional Shoc2 or expressing Shoc2 NSLH variants.

To further assess the role of PAK1 and 2 in the phosphorylation of ERK1/2 in cells expressing Shoc2 variants, we used a small-molecule PAK1 and 2 inhibitor, FRAX1036 (47). Cells were treated with 2.5 µM FRAX1036 for 24 hours and then stimulated with 30 ng/ml of EGF (**Fig. 5**). Similar to the experiments in **Fig. 4**, we found that EGF treatment resulted in increased phosphorylation of PAK in cells expressing the Shoc2^S2G^ or Shoc2^M173I^ variants (**Fig. 5A, B and C**). FRAX1036 treatment abolished the phosphorylation of PAK and dramatically reduced levels of ERK1/2 phosphorylation in cells expressing Shoc2^S2G^ or Shoc2 ^M173I^ variants. FRAX1036 treatment had little effect on ERK1/2 phosphorylation in cells expressing WT Shoc2 (**Fig. 5A, B, and D**). These data suggest that PAKs contribute to increased pERK1/2 phosphorylation in cells lacking functional Shoc2. Thus, we conclude that high EGF concentration-induced activation of the AKT-PAK feedback loop enhances ERK1/2 signaling in cells expressing Shoc2 variants. The observations that low EGF concentrations do not activate the AKT-PAK axis supported this conclusion (**Supp.** Fig. 3).

**Figure 5.**
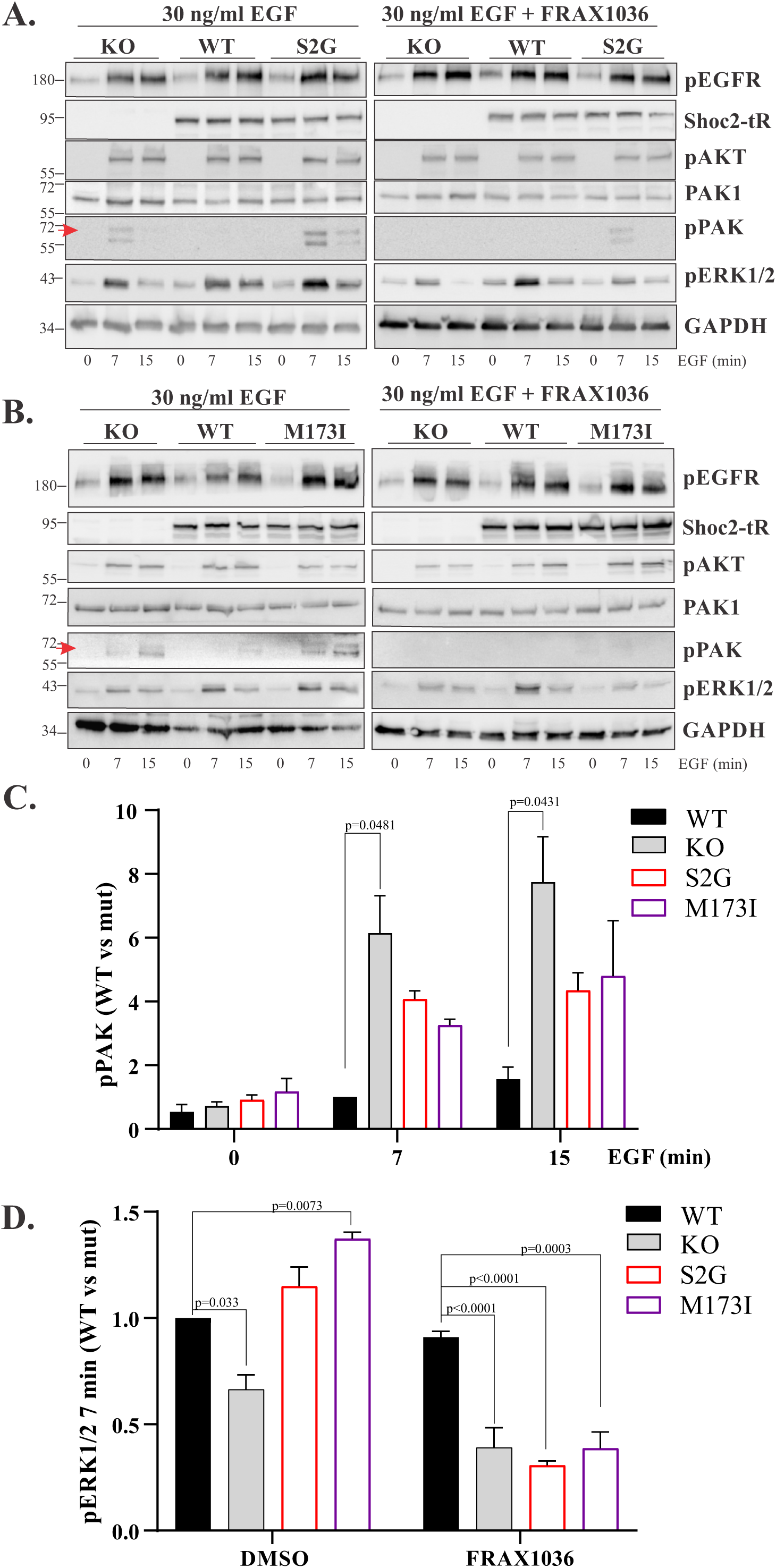
High EGF concentrations activate the AKT-PAK axis in a Shoc2-independent manner. **A.** HeLa-KO cells transiently transfected with the Shoc2-tRFP or Shoc2-tRFP (S2G) mutant were serum-starved and treated with DMSO or FRAX1036 for 24 hr and then stimulated with EGF (0.2 and 30 ng/ml) for 7-or 15-min at 37°C. Cell lysates were analyzed using anti-pEGFR, –Shoc2, –pERK1/2, –pAKT, –pPAK, –PAK1, AKT, and –GAPDH antibodies. **B.** HeLa-KO cells transiently transfected with the Shoc2-tRFP, Shoc2-tRFP (M173I) mutant were serum-starved and treated with DMSO or FRAX1036 for 24 hr, and then stimulated with EGF (0.2 and 30 ng/ml) for 7-or 15-min at 37^°^C. Cell lysates were analyzed using anti-pEGFR, – Shoc2, –pERK1/2, –pAKT, –pPAK, AKT, –PAK1, and –GAPDH antibodies. **C.** Bars represent the mean amount of pPAK in cells not treated with FRAX1036 and expressing Shoc2 variants normalized to the total amount of GAPDH in arbitrary units as compared to cells expressing WT Shoc2-tRFP at 7min ± S.E. (*n*=3) *(p values as shown on graph,* Anova one way). The results in each panel are representative of those from three independent experiments. **D.** Bars represent the mean amount of pERK1/2 in cells expressing Shoc2 variants normalized to the total amount of GAPDH in arbitrary units as compared to cells expressing WT Shoc2-tRFP at 7min ± S.E not treated with FRAX1036. (*n*=3) *(p values shown on the graph,* Anova one way). The results in each panel are representative of those from three independent experiments. Red arrows denote the pPAK chemiluminescence signal used for quantification.

As noted above, PAK1 stimulates the phosphorylation of S338 of RAF1 and S298 of MEK1/2 in response to EGF, thereby priming and activating MEK1/2 for phosphorylation of ERK1/2 (T202/Y204) (45,48,49). Hence, we dissected molecular events further and examined how FRAX1036 treatment affects the corresponding phosphorylation of RAF1 and MEK1/2 kinases. HeLa Shoc2 KO cells expressing Shoc2^WT^ or the Shoc2^S2G^ variants were treated with 2.5 µM FRAX1036 for 24 hours and then stimulated with 30 ng/ml of EGF (**Fig. 6**). We found that FRAX1036 treatment reduced the phosphorylation of RAF1 and MEK1/2 in cells expressing Shoc2^S2G^ Shoc2 variant. RAF1 phosphorylation in cells expressing Shoc2 WT remained largely unaffected. Data in **Fig. 6** support our conclusions that PAKs contribute to increased ERK1/2 phosphorylation in cells lacking functional Shoc2. We report herein that, in cells expressing Shoc2 NSLH variants or lacking Shoc2, high EGF concentrations trigger the AKT-PAK-ERK1/2 signaling loop.

**Figure 6.**
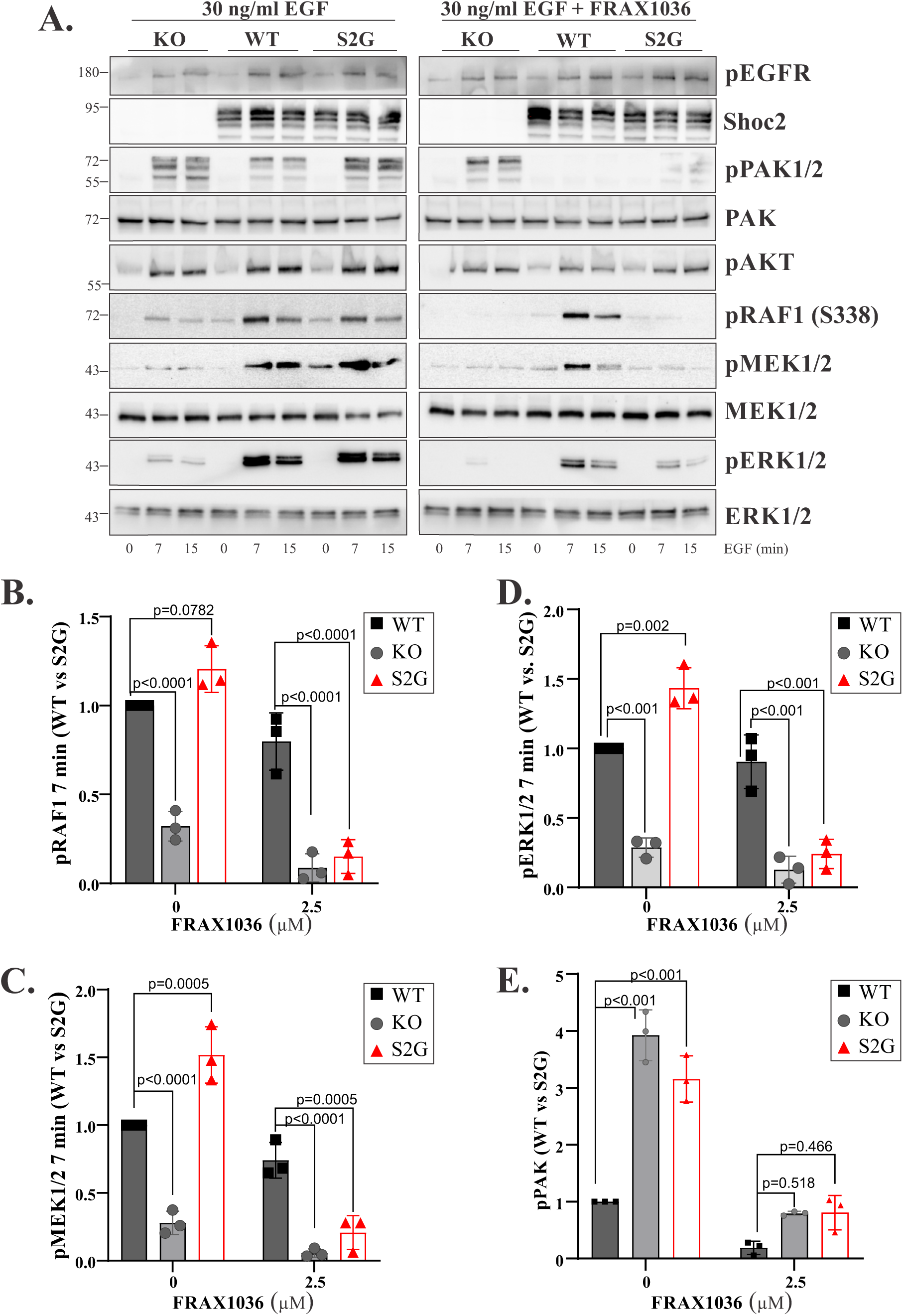
The AKT-PAK signaling feedback loop stimulates RAF1 and MEK1/2 phosphorylation in cells activated with high ligand concentrations. **A.** HeLa-KO cells transiently transfected with the Shoc2-tRFP or Shoc2-tRFP (S2G) mutant were serum-starved and treated with DMSO or FRAX1036 for 24 hr and then stimulated with EGF (30 ng/ml) for 7-or 15-min at 37°C. Cell lysates were analyzed using anti-pEGFR, –Shoc2, –pERK1/2, –pAKT, –pPAK, –PAK1, AKT, RAF1, MEK1/2 and –GAPDH antibodies. **B-E.** Bars represent the mean amount of pRAF1 (**B**), pMEK1/2 (**C**), pERK1/2 (**D**) and pPAK (**E**) in cells not treated with FRAX1036 and expressing Shoc2 variants normalized to the total amount of ERK in arbitrary units as compared to cells expressing WT Shoc2-tRFP at 7min ± S.D. (*n*=3) *(p values as shown on graph,* two-way ANOVA). The results in each panel are representative of those from three independent experiments.

Our finding that high EGF concentrations induce AKT/PAK-mediated phosphorylation of ERK1/2 in cells lacking the functional Shoc2 scaffold delivered a critical clue to the model in **Fig. 7**. In this model, under the conditions when Shoc2 guides EGF signal primarily through the Ras-ERK1/2 pathway, Shoc2’s genetic NSLH variants cannot maintain the amplitude of ERK1/2 signals. Conversely, when triggered, the AKT-PAK signaling feedback can activate the ERK1/2 pathway in a Shoc2-independent manner. Treatment with either AKT or PAK inhibitors prevented the feedback activation of ERK1/2. Hence, our studies identified a new regulatory AKT-dependent feedback loop activated in the presence of the Shoc2 NSLH variants.

**Figure 7.**
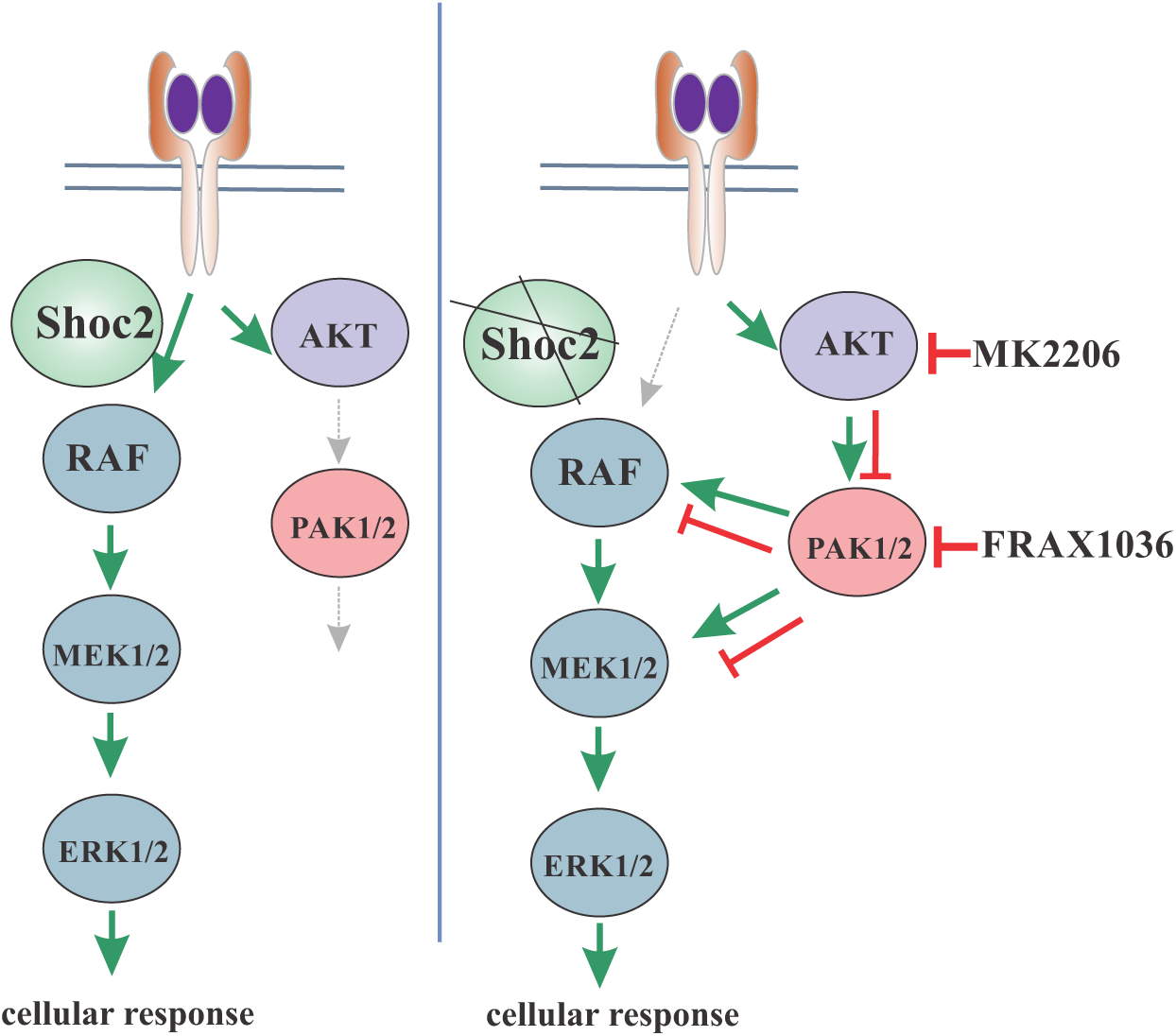
Summary depicting the crosstalk between the Shoc2-mediated AKT and ERK1/2 signals. Model showing the interplay between the ERK1/2 and AKT signaling pathways in the presence and absence of the functional Shoc2 when cells are stimulated with high EGF concentrations. In cells either lacking Shoc2 or expressing Shoc2 pathogenic variants, AKT activates PAK1-2 kinases, thereby inducing Shoc2-independent phosphorylation of ERK1/2.

## DISCUSSION

The present study evaluates signaling outcomes downstream of EGFR in cells expressing genetic variants of the non-enzymatic scaffold protein Shoc2. As reported earlier, EGF concentrations as low as 0.2 ng/ml, ∼34 pM, can stimulate nearly 80% of ERK1/2 phosphorylation. In comparison, EGF concentrations such as 20-30 ng/ml, ∼5 nM) also activate the AKT pathway (**Supp.** Fig. 1). These findings are well-aligned with the work of others demonstrating that activation of the small pool of the high-affinity EGFR is sufficient to stimulate maximal ERK1/2 phosphorylation (33–36). Our study shows that Shoc2 loss abrogates ERK1/2 phosphorylation but does not affect the activation of AKT. Furthermore, activation of the AKT pathway was unaffected by the Shoc2 NSLH mutations. Significantly, our experiments demonstrate that activating different signaling pathways downstream of EGFR impacts ERK1/2 phosphorylation in cells expressing NSLH variants, emphasizing the importance of “contextual” signaling (**Fig. 2 and 3**).

The analysis of ERK1/2 and the AKT intracellular signals in the context of Shoc2 NSLH mutations allowed us to identify a previously unrecognized AKT-mediated feedback loop. Data in **Fig. 4 and 5** shows that in cells expressing Shoc2 NSLH-causing variants or lacking Shoc2, the AKT pathway mediates phosphorylation of the p21-activated (PAK) kinases 1 and 2. We found that inhibiting PAK1 and 2 using FRAX1036 inhibitor abolishes RAF1(S338) and MEK1/2 or phosphorylation (**Fig. 6**). Together, these data indicate that PAK-mediated feedback loop contributes to increased ERK1/2 activation in cells expressing Shoc2 NSLH mutants. Both PAK family kinases are critical downstream effectors of EGFR (48,50,51), and PAK1 was shown to activate the ERK1/2 pathway in response to EGF (45,48). Several studies have demonstrated that PAK1 stimulates the phosphorylation of S338 of RAF1 and S298 of MEK1/2, thus priming and activating MEK1/2 for phosphorylation of ERK1/2 (T202/Y204) (45,46,49).

Interestingly, PAK1 was also implicated in the regulation of ERK1/2 as a scaffold that recruits MEK to RAF-1 at the plasma membrane (52), thus facilitating signal transmission through the ERK1/2 pathway (53,54). We have not examined whether 30ng/ml of EGF triggers the activation of phosphatidylinositol 3-kinase (PI3K), followed by the activation of Rac, AKT, and, subsequently, PAK1 (55). Yet, this would be consistent with prior reports demonstrating the EGF-dependent Shoc2 interaction with the p110α subunit of PI3K, Ras, and RAF-1 in cells stimulated with 20 ng/ml of EGF (13).

Notably, our data show that in cells expressing WT Shoc2 and exposed to high EGF concentrations, levels of phosphorylated PAK1 and 2 were negligible (**Fig. 4 and 5**), highlighting the critical role of Shoc2 in preferentially transmitting the EGFR-ERK1/2 signals and further supporting the notion of Shoc2 as a “gatekeeper.”

At least seven different ligands can activate human EGFR and generate signals to several downstream pathways, including ERK1/2, phosphoinositide 3-kinase (PI3K), AKT, Src family kinases (SFKs), STATs, phospholipase Cγ1, and others (7,56–58). Many of these signaling cascades are interconnected and merge into an EGFR-stimulated signaling network – a subject covered by a large body of literature. The signaling outputs of EGFR depend on various factors, and each EGFR ligand elicits qualitatively and quantitatively different downstream signals (33,59,60). Furthermore, in the context of the specific ligand, signaling responses downstream of EGFR depend on the number of activated receptors (i.e., signal intensity) (37,61,62) and the distinct affinities with which EGFRs bind to its ligands (35,63,64). It is well-established that stimulation of the large pool of EGF receptors contributes to the parallel activation of several signaling pathways, including the ERK1/2 and AKT cascades (36). Therefore, assuming that WT Shoc2 transduces signals of the small pool of high-affinity EGFRs is plausible.

NSLH is one of several developmental disorders caused by germline mutations in genes encoding protein components of the Ras/ERK1/2 pathway (65). When the pathogenicity of newly identified patient mutations is determined, the levels of phospho-ERK1/2 downstream of EGFR are employed as an indicator for enhanced protein activity (e.g., gain-of-function) or a loss of its native function (e.g., loss-of-function). In most of these studies, cells are stimulated with EGF concentrations within the 20–30 ng/ml range. For proteins such as PTPN11 or Ras, for which secondary examination is possible, assays directly assessing phosphatase activity of purified proteins *in vitro* or analysis of the intrinsic and guanine nucleotide exchange factor – accelerated nucleotide exchange can confirm gain-of-function or loss-of-function nature of mutations (66,67). However, for proteins without apparent enzymatic activity, including the Shoc2 scaffold, such assays are not available. Thus, other factors, such as using EGF concentrations relevant to the concentrations found in the human body, are significant. The EGF concentrations recorded in amniotic fluid and fetal urine are in pg/ml levels and remain low during the early neonatal period (68). Although these levels rise in adult tissue, they range from ∼0.2-0.6 ng/ml in serum to ∼60-80 ng/ml in urine (29,31,69,70). In addition, it is well-established that EGF doses as low as 2 pM activate the ERK1/2 signaling cascade *in vivo* (64) and that signaling outcomes observed in response to picomolar EGF concentrations versus nanomolar EGF concentrations activate distinct signaling pathways in cultured cells (35,36,71–73). Hence, for assessing protein variants for which secondary assay is unavailable, determinants such as activation of the specific downstream pathway under physiological conditions become critical. On note, our zebrafish model studies showed that embryos injected with wild-type *shoc2* mRNA rescued the shoc2 morpholino-induced defects in erythrocyte circulation. In contrast, in embryos injected with *shoc2* mRNA harboring Shoc2^S2G^, Shoc2^L437I,^ or Shoc2^QH269/270YH^ mutations, erythropoiesis was not rescued to the level of wild-type Shoc2 (15), further supporting the importance of assessing pathogenic mutations under the conditions close to physiological.

The studies resolving the Shoc2-MRas-PP1C complex structure suggested that the Shoc2 variant QH269/270HY potentially forms more extensive contacts between PP1CA and may increase the affinity of the protein in the complex. It was also suggested that the M173I variant might create a de novo contact with MRAS, possibly extending the lifetime of the Shoc2-MRas-PP1C complex and leading to sustained RAF dephosphorylation. Yet, the experimental conditions assessing the complex assembly in the presence of the activated RAS (GTP-bound RAS mutant) do not account for the intricate intracellular events and a complex network of signaling crosstalk induced by the “contextual” signaling events. It also remains unclear how the NSLH-related mutations affect Shoc2 interactions with other Ras isoforms (H-, K-, and N-RAS), which are likely to be formed when low numbers of EGFR are activated. These and other questions will be resolved in future studies using different models.

The role of Shoc2 in cancer progression has been explored as well. Even though there is not enough evidence to suggest that NSLH Shoc2 mutations cause pediatric cancers, instances of neuroblastoma and cutaneous T-cell lymphoma have been reported in patients heterozygous for the c.4A>G Shoc2 variant (5,74). Several studies have suggested that Shoc2 could be an independent prognostic marker for breast, lung, and other cancers (75) or have evaluated Shoc2’s feasibility as a therapeutic target for pancreatic, lung, and colorectal tumors (76–78). Moreover, Shoc2 has been explored as a target for sensitization of the MEK inhibitors in non-small-cell lung cancers (77). Somatic Shoc2 mutations have been found in leukemia biopsies, glioblastomas, and urinary cancers (78–80). The relevance of our findings to the progression of different tumors remains to be established, and future investigations addressing the role of somatic Shoc2 mutations in tumorigenesis should assess the AKT-PAK signaling axis.

In summary, by analyzing signaling responses guided by the Shoc2 scaffold in the context of different signaling cascades, we show that EGF concentrations, the number of activated receptors, and possible feedback loop mechanisms should be considered when functionally assessing Shoc2’s disease-causing genetic variants. Yet, to fully appreciate the physiological relevance of our findings for the molecular diagnostics of patients with NSLH, new animal models and/or patient samples are necessary. The most accurate understanding of the context in which signals are transmitted will yield a clearer picture of how Shoc2 NSLH genetic variants affect the function of the scaffold. A better grasp of contextual signaling will improve our knowledge of developmental pathologies, help predict cellular outcomes when evaluating the effects of hereditary Shoc2 variants on different tissues, and lead to the design of more effective and individualized therapies for RASopathies and cancers.

## Materials and Methods

### Antibodies and other reagents

Specific proteins were detected using primary antibodies to the following: phosphorylated ERK1/2 (pERK1/2), Shoc2 and GAPDH (Santa Cruz Biotechnology, Dallas, TX, USA); phosphorylated –EGFR (Y1068), phosphorylated –AKT (S473), phosphorylated – PAK1 (Thr423)/PAK2 (Thr402) (Cell Signaling, Danvers, MA, USA); total PAK1 and total AKT (Proteintech, Rosemont, IL, USA); Peroxidase-conjugated AffiniPure F(ab’)2 Fragment Goat Anti-Rabbit and –Mouse IgG (H+L) (Jackson Immuno). EGF was purchased from BD Biosciences, MK2206 from Tocris Bioscience, and FRAX1036 from Selleckchem (Huston, TX, USA).

### Expression plasmids

tRFP-tagged Shoc2 (Shoc2-tRFP) was described previously (81). Shoc2-tRFP-tagged mutants were generated as described previously (81) and (15). All constructs were verified by dideoxynucleotide sequencing.

### Cell culture and DNA transfections

293FT cells (Invitrogen), Cos1cells (ATCC), HeLa cells (ATCC), and stable cell lines (derivatives of HeLa cells) were grown in Dulbecco Modified Eagle’s Medium (DMEM) containing 10% fetal bovine serum (FBS) supplemented with sodium pyruvate, minimal essential medium with nonessential amino acids (MEM-NEAA), penicillin, streptomycin, and L-glutamate (Invitrogen). Transfections of DNA constructs were performed using the polyethyleneimine (Neo Transduction Laboratories, Lexington, KY) or Jetoptimus (Polyplus LLC) reagent. The expression of proteins was confirmed by Western blotting, as described below.

### CRISPR/Cas9-mediated gene deletion

A HeLa cell line lacking Shoc2 was generated by Biocytogen (http://www.biocytogen.com). RNA guide sequences (5’GATAAAGGTATTGCCTCTGT TGG3’ and 5’GGAATAAAGGTCAAAAGATT AGG3’) targeting exon 2 and Cas9/CRISPR technology were used to insert a puromycin resistance gene. Clones were isolated, screened genetically, and then tested for Shoc2 expression by immunoblot. Gene disruption was also validated by PCR analysis.

### Western blot analysis

Cells were placed on ice and washed with Ca^2+^, Mg^2+^-free phosphate buffered saline (PBS), and the proteins were solubilized in 50 mM Tris (pH 7.5) containing 150 mM NaCl, 1% Triton X-100, 1 mM Na_3_VO_4_, 10 mM NaF, 0.5 mM phenylmethylsulfonyl fluoride (PMSF, Sigma, St. Louis, MO, USA), 10 μg/ml of leupeptin, and 10 μg/ml of aprotinin (Roche, Basel, Switzerland) for 15 min at 4°C. Lysates were centrifuged at 14,000 *rpm* for 15 min to remove insoluble material. Aliquots of cell lysates were denatured in sample buffer at 95°C, resolved by electrophoresis, and probed by Western blotting with various antibodies followed by chemiluminescence detection.

For western blot analysis of zebrafish larvae, proteins were extracted from dechorionated and de-yolked embryos into a buffer containing 150mM NaCl, 1% Triton-X 100, 0.5% Sodium deoxycholate, 0.1% SDS, 50mM Tris pH 8.0, 1 mM Na_3_VO_4_, 10 mM NaF, 0.5 mM PMSF, 10 μg/ml of leupeptin, and 10 μg/ml of aprotinin. Western blotting was done as described previously (81). Proteins transferred from SDS-polyacrylamide gels to nitrocellulose membranes were visualized using a ChemiDoc analysis system (Bio-Rad, Hercules, CA, USA). Several exposures were analyzed to determine the linear range of the chemiluminescence signals. Quantification was performed using the densitometry analysis mode of Image Lab software (Bio-Rad, Hercules, CA, USA). To visualize proteins with close molecular weight, in some instances, two identical SDS-PAGE gels were used to resolve protein lysate.

### Zebrafish strains and maintenance

All zebrafish strains were bred, raised, and maintained following established animal care protocols for zebrafish husbandry. Embryos were staged as previously described (82). All animal procedures were carried out by guidelines established by the University of Kentucky Institutional Animal Care and Use Committee.

### Morpholino and mRNA injection

All MOs were obtained from Gene Tools, LLC (Philomath, OR) and injected into 1-2 cell stage zebrafish embryos. The following MOs were used in this study: standard control MO: 5’-CCTCTTACCTCAGTTACAATTTATA-3’; *shoc2* MO: 5’-TACTGCTCATGGCGAAAGCCCCGCA-3’. Embryos were injected with 8 ng each of MOs. For mRNA rescue experiments, the zebrafish *shoc2* coding sequences (with silent mutations at MO target sites) for either WT or mutant were PCR amplified from zebrafish *shoc2* complementary DNA (cDNA) and cloned into the pCS2+ vector (Promega, WI). The capped mRNAs were synthesized with the mMessage mMACHINE SP6 transcription kit (Thermofisher Scientific, CA) according to the manufacturer’s instructions. For mRNA rescue experiments, 100 pg/embryo of WT was co-injected with 8ng of *shoc2* MO.

### Statistical analyses

Results are expressed as means ± S.D. The statistical significance of the differences between groups was determined using either Student’s *t*-test or one-way and two-way ANOVA (followed by Tukey’s test or Kruskal-Wallis test). *p* < 0.05 was considered statistically significant. GraphPad Prism 8.0 (GraphPad Prism Software Inc., Chicago, IL, USA) was used to perform all statistical analyses.

### Data availability

This published article and its supporting information files include all data generated or analyzed during this study.

### Supporting information

This article contains supporting information.

## Acknowledgments

We thank Drs. Charles Waechter and Louis Hersh for critical reading of the manuscript.

## Funding

This project was supported by grants from the National Institute of General Medical Sciences (R35GM136295 and 1S10OD025033-01 to EG). Its contents are solely the authors’ responsibility and do not necessarily represent the official views of the National Institute of Health.

***Conflict of interest statement:*** none declared

## SUPPORTING FIGURE LEGENDS

**Supporting Figure 1.** The ERK1/2 pathway is the primary signaling cascade activated by low ligand concentrations. **A.** serum-starved HeLa cells were incubated with 0-32 ng/ml EGF for 7 min at 37°C. Cell lysates were probed with anti-pEGFR, –pERK1/2, –pAKT, and –GAPDH antibodies. Numbers indicate the mean amount of pERK1/2 normalized to the total amount of GAPDH in arbitrary units compared to cells treated with 32ng/ml at 7min ± S.D. (*n*=3). The results in each panel are representative of those from three independent experiments.

**Supporting Figure 2.** The effect of Shoc2 NSLH mutant on ERK1/2 phosphorylation in cells expressing endogenous Shoc2. **A**. HeLa cells transiently transfected with the Shoc2-tRFP were serum-starved for 16 hr and then stimulated with EGF (0.2 ng/ml) for 7-or 15-min at 37°C. Cell lysates were analyzed using anti-pEGFR, –pERK1/2, –ERK1/2, –pMEK1/2, –GAPDH, and –Shoc2 antibodies. Bars represent the mean amount of pERK1/2 in HeLa cells normalized to the total amount of GAPDH in arbitrary units as compared to cells expressing WT Shoc2-tRFP at 7min ± S.E. (*n*=3) *(p values as shown on graph,* two-way ANOVA). The results in each panel are representative of those from three independent experiments. **B.** Cos1 cells transiently transfected with the Shoc2-tRFP or Shoc2-tRFP S2G mutant were serum-starved for 16 hr and then stimulated with EGF (0.2 and 30 ng/ml) for 7-or 15-min at 37°C. Cell lysates were analyzed using anti-pEGFR, –pERK1/2, –pAKT, –GAPDH, and –Shoc2 antibodies. **C.** Bars represent the mean amount of pERK1/2 in Cos1 cells expressing Shoc2 mutants normalized to the total amount of GAPDH in arbitrary units as compared to cells expressing WT Shoc2-tRFP at 7min ± S.D. (*n*=4) *(p values as shown on graph,* two-way ANOVA). The results in each panel are representative of those from three independent experiments. **D.** 293FT cells transiently transfected with the Shoc2-tRFP or Shoc2-tRFP S2G mutant were serum-starved for 16 hr and then stimulated with EGF (0.2 and 30 ng/ml) for 7-or 15-min at 37°C. Cell lysates were analyzed using anti-pEGFR, –pERK1/2, –pAKT, –GAPDH, and –Shoc2 antibodies. **E.** Bars represent the mean amount of pERK1/2 in 293FT cells expressing Shoc2 mutants normalized to the total amount of GAPDH in arbitrary units as compared to cells expressing WT Shoc2-tRFP at 7min ± S.D. (*n*=4) *(p values as shown on the graph,* two-way ANOVA). The results in each panel are representative of those from three independent experiments.

**Supporting Figure 3.** Low EGF concentrations do not activate the AKT-PAK axis in a Shoc2-independent manner. **A.** HeLa Shoc2 CRISPR KO cells transiently transfected with the Shoc2-tRFP or Shoc2-tRFP S2G mutant were serum-starved for 16 hr and then stimulated with EGF (0.2 ng/ml) for seven or 15min at 37°C. Cell lysates were analyzed using anti-pEGFR, –pERK1/2, –pAKT, –AKT, –pPAK, –PAK1, –GAPDH and –Shoc2 antibodies.

## Notes

### Competing Interest Statement

The authors have declared no competing interest.

