## Supplementary figures and images for "The expression of congenital Shoc2 variants induces AKT-dependent feedback activation of the ERK1/2 pathway"

### Supplemental figures 1-3

#### Supporting Figure 1

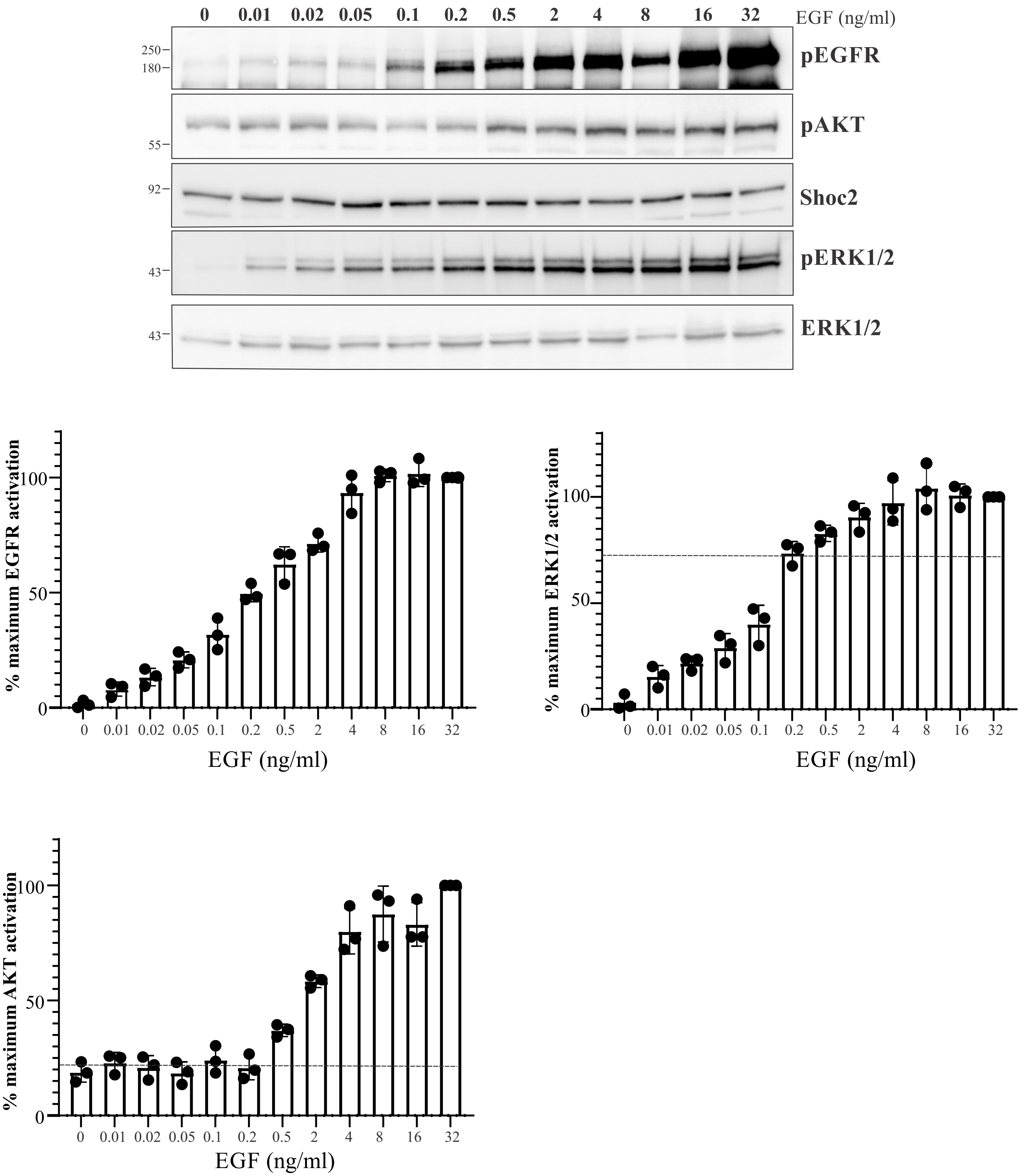

#### Supporting Figure 2

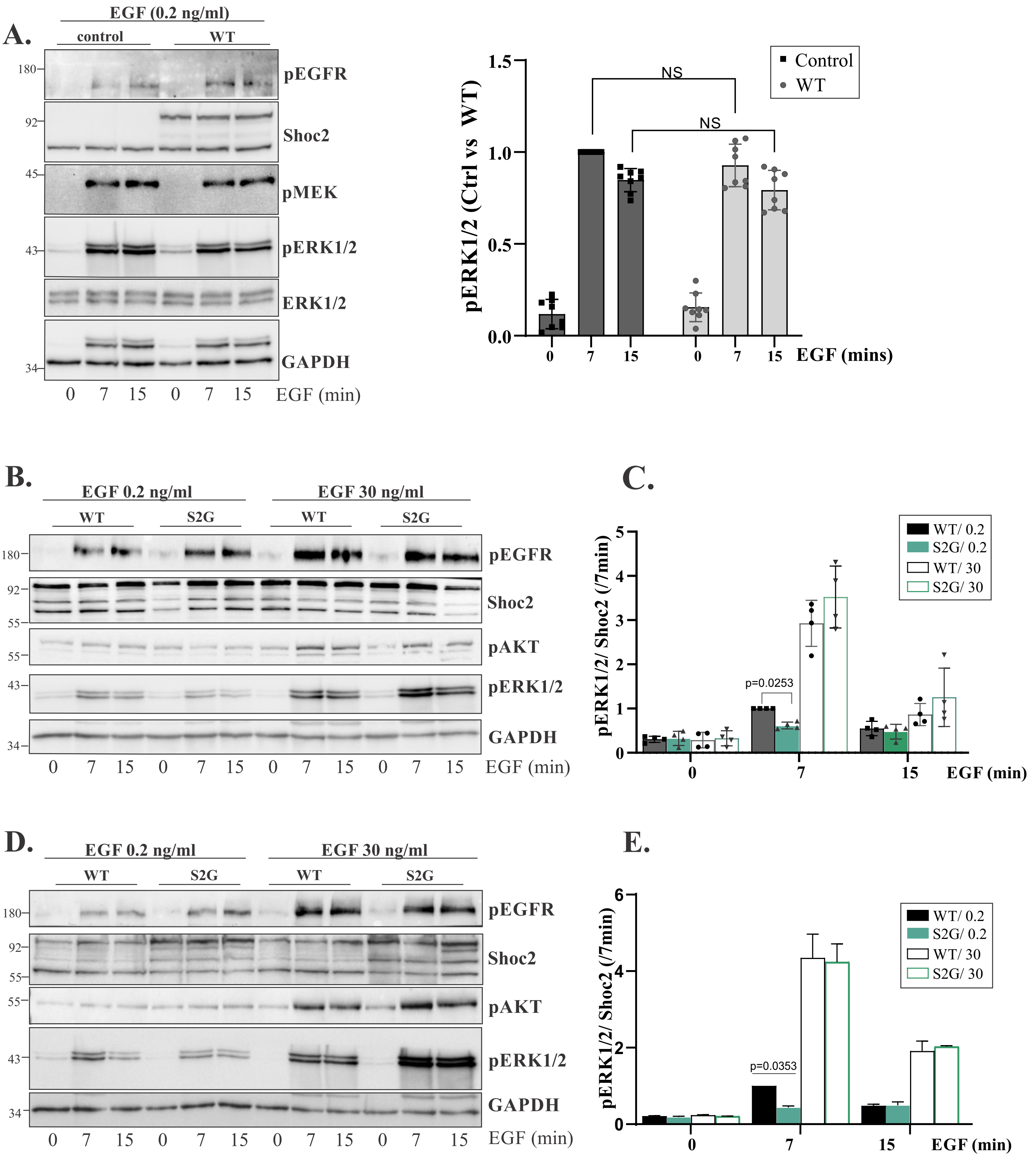

##### Supporting Figure 3

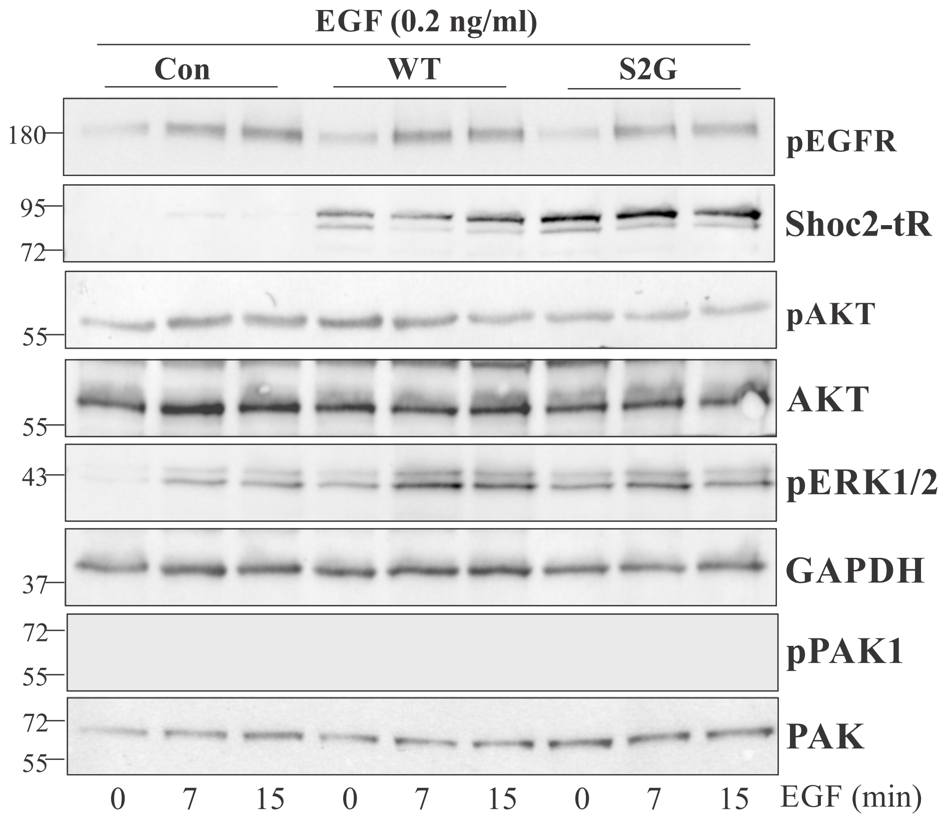
